# CO_2_ conversion to methane and biomass in obligate methylotrophic methanogens in marine sediments

**DOI:** 10.1101/528562

**Authors:** Xiuran Yin, Weichao Wu, Mara Maeke, Tim Richter-Heitmann, Ajinkya C. Kulkarni, Oluwatobi E. Oni, Jenny Wendt, Marcus Elvert, Michael W. Friedrich

**Author notes:** Correspondence: Michael W. Friedrich, Microbial Ecophysiology Group, Faculty of Biology/Chemistry, University of Bremen, PO Box 33 04 40, D-28334 Bremen, Germany. These authors contributed equally to this work (Xiuran Yin, Weichao Wu). Current address: Department of Biogeochemistry of Agroecosystems, University of Goettingen, Goettingen, Germany (Weichao Wu).

## Abstract

Methyl substrates are important compounds for methanogenesis in marine sediments but diversity and carbon utilization by methylotrophic methanogenic archaea have not been clarified. Here, we demonstrate that RNA-stable isotope probing (SIP) requires ^13^C-labeled bicarbonate as co-substrate for identification of methylotrophic methanogens in sediment samples of the Helgoland mud area, North Sea. Using lipid-SIP, we found that methylotrophic methanogens incorporate 60 to 86% of dissolved inorganic carbon (DIC) into lipids, and thus considerably more than what can be predicted from known metabolic pathways (∼40% contribution). In slurry experiments amended with the marine methylotroph *Methanococcoides methylutens*, up to 12% of methane was produced from CO_2_, indicating that CO_2_-dependent methanogenesis is an alternative methanogenic pathway and suggesting that obligate methylotrophic methanogens grow in fact mixotrophically on methyl compounds and DIC. Thus, the observed high DIC incorporation into lipds is likely linked to CO_2_-dependent methanogenesis, which was triggered when methane production rates were low. Since methylotrophic methanogenesis rates are much lower in marine sediments than under optimal conditions in pure culture, CO_2_ conversion to methane is an important but previously overlooked methanogenic process in sediments for methylotrophic methanogens.

## Introduction

Methanogenesis is the terminal step of organic matter mineralization in marine sediments [1]. There are three main pathways producing methane, i.e., hydrogenotrophic (H_2_/CO_2_), acetoclastic (acetate) and methylotrophic (e.g., methanol, methylamine) methanogenesis [2] with the former two pathways considered dominant [3]. However, the importance of methylated compounds for methanogenesis in marine sediments has been acknowledged in recent years. Geochemical profiles and molecular analysis have shown that methylotrophic methanogenesis is the most significant pathway for methane formation in hypersaline sediments [4,5] and in the sulfate reduction zone (SRZ) in marine environments [6,7] where methanol concentration of up to 69 µM had been measured [8,9]. Especially in the SRZ, methylated compounds are regarded as non-competitive substrates for methanogenesis, since sulfate reducing microorganisms apparently do not compete with methanogens for these compounds [10]. In sediments of the Helgoland mud area, specifically, high relative abundances of potential methylotrophic methanogens were observed [11] of which many are unknown. The potential for methylotrophic methanogenesis was recently even predicted from two metagenome-assembled genomes of uncultivated Bathyarchaeota, assembled from a shotgun metagenome [12].

The formation of methane via the three main pathways in methanogenic archaea has been studied intensively [13,14,15]. Much less is known regarding assimilation of carbon into biomass under *in situ* conditions and to which extend different carbon sources in the environment are utilized. Discrepancies between predicted pathways known and the actual carbon metabolism measured appear to be based on different cellular functions of carbon dissimilation and assimilation, originate from reaction equilibria operative, intermediate carbon cross utilization as well as interplay between different microbial communities [14, 16,17,18,19]. For example, mixotrophically growing cultures of *Methanosarcina barkeri* form their biomass equally from methanol and CO_2_, however, almost all the methane is formed from methanol rather than from CO_2_ [20] since methanol is disproportionated to methane and CO_2_ according to the following reaction:

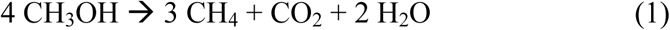

But apart from such culture studies using the nutritionally versatile *Methanosarcina barkeri*, which is capable of hydrogenotrophic, acetoclastic and methylotrophic methanogenesis, the respective contribution of CO_2_ and methylated carbon substrates to biomass formation during methylotrophic methanogenesis, especially for “obligate” methylotrophic methanogens, in natural sediments has not been studied to date.

Nucleic acids (RNA ∼20%, DNA ∼3%, of dry biomass, respectively), lipids (7–9%) and proteins (50–55%) are crucial cell components in living microorganisms [21], and thus, are suitable markers of carbon assimilation. In order to characterize carbon assimilation capabilities, stable isotope probing (SIP) techniques exist, among which RNA-SIP is very powerful for identifying active microorganisms based on separating ^13^C-labeled from unlabeled RNA using isopycnic centrifugation [22,23]. In combination with downstream sequencing analysis, RNA-SIP provides high phylogenetic resolution in detecting transcriptionally active microbes [24,25] but is limited in its sensitivity by requiring more than 10% of ^13^C incorporation into RNA molecules for sufficient density gradient separation [26]. To date, a number of SIP studies successfully detected methylotrophic bacteria [27,28,29] but the detection of methylotrophic methanogens by RNA-SIP with ^13^C labeled methyl compounds might be hampered by mixotrophic growth [20], i.e., simultaneous assimilation of carbon from methylated compounds and CO_2_.

In contrast to RNA-SIP, lipid-SIP has a lower phylogenetic resolution, but can detect very sensitively δ^13^C-values in lipid derivatives by gas chromatography combustion isotope ratio mass spectrometry (GC-IRMS), thereby facilitating quantitative determination of small amounts of assimilated carbon [30,31].

In this study, we aimed to identify methylotrophic methanogens by RNA-SIP and elucidate carbon assimilation patterns in marine sediments. We hypothesized that the large pool of ambient dissolved inorganic carbon (DIC) in sediments [6] alters carbon utilization patterns in methylotrophic methanogens compared to pure cultures. To address these questions, we tracked carbon dissimilation into methane and quantified assimilation into lipids by lipid-SIP in slurry incubations and pure cultures. We found an unexpectedly high degree of methanogenesis from DIC by previously considered obligate methylotrophic methanogens, i.e., using only methyl groups for methane formation. This mixotrophic methanogenesis might be the basis for our observation that more inorganic carbon was assimilated into biomass than could be expected from known pathways.

## Materials and Methods

### Sediment incubation setup for SIP

Sediment was collected from the Helgoland mud area (54°05.23’N, 007°58.04’E) by gravity coring in 2015 during the RV HEINCKE cruise HE443. The geochemical profiles were previously described [11]. Sediments of the SRZ (16–41 cm) and MZ (238–263 cm) from gravity core HE443/077-1 were selected for incubations. 50 mL of 1:4 (w/v) slurries were prepared anoxically by mixing sediments with sterilized artificial sea water without sulfate (26.4 g L^−1^ NaCl, 11.2 g L^−1^ MgCl_2_·6H_2_O, 1.5 g L^−1^ CaCl_2_·2H_2_O and 0.7 g L^−1^ KCl). Slurries were dispensed into sterile 120-mL serum bottles and sealed with butyl rubber stoppers. Residual oxygen was removed by exchanging bottle headspace 3 times with N_2_ gas. A 10-day pre-incubation was performed, followed by applying vacuum (3 min at 100 mbar) to remove most of the headspace CO_2_. Triplicate incubations were conducted by supplementing 1 mM ^13^C-labeled methanol and unlabeled 10 mM sodium bicarbonate, or 1 mM unlabeled methanol and 10 mM^13^C-labeled sodium bicarbonate (^13^C-labeled substrates provided by Cambridge Isotope Laboratories, Tewksbury, Massachusetts, USA) at 10 °C.

### Pure culture setup

The carbon assimilation patterns were compared between SIP sediment incubations and the obligate methylotrophic methanogen, *Methanococcoides methylutens*. *M. methylutens* strain MM1 (DSM 16625) was obtained from the German Collection of Microorganisms and Cell Cultures (DSMZ, Braunschweig, Germany). Initial cultivation was performed using Medium 280 according to DSMZ protocols. After several transfers of the culture in anoxic marine Widdel medium [32], 5% of the culture were inoculated into fresh Widdel medium supplemented with 30 mM methanol, trace element solution SL 10 [33], and 50 mM sodium bicarbonate (i.e., DIC) with carbon sources containing 5% of ^13^C-label. Pure cultures were grown at 30 °C. Cells were harvested after methane formation had ceased by centrifugation and used for lipid-SIP analysis.

### Slurry incubations inoculated with *M. methylutens*

To test methanogenesis from CO_2_, a series of incubations were performed with *M. methylutens* in autoclaved (n=3) sediment slurry from the SRZ with different amendments of electron donor (H_2_), electron shuttles (humic acid; anthraquinone-2,6-disulfonic acid - AQDS), and electron acceptors/electron conductors (hematite, α-Fe_2_O_3_; magnetite, Fe_3_O_4_; Lanxess, Germany). Incubations were separately prepared with 50% H_2_ in headspace, 100 µM AQDS, 30 mM magnetite, 30 mM hematite and 500 mg L^−1^ humic acid (Sigma-Aldrich, Steinheim, Germany). 5% of the culture was inoculated into these setups each, and amended with 20 mM unlabeled methanol and 10% of ^13^C-labeled DIC for measuring carbon partitioning into methane. The control incubation comprised autoclaved slurry, 50% H_2_ in headspace and 20 mM methanol without addition of *M. methylutens*. All experiments were setup with a total volume of 50 mL in 120-mL serum bottles sealed with butyl rubber stoppers, and incubated at 30?°C.

### Gas analysis

The concentration of methane in the headspace of bottles was measured by gas chromatography as previously described [34]. Headspace H_2_ in incubation bottles was determined with a reduction gas detector (Trace Analytical, Menlo Park, California, USA). 1 mL was used for injection. The parameters were as follows: carrier gas (nitrogen) 50 mL min^−1^, injector temperature 110 °C, detector 230 °C, column (Porapak Q 80/100) 40 °C.

The δ^13^C values of methane and CO_2_ in the headspace as well as of DIC in the slurries were determined using a Thermo Finnigan Trace GC connected to a DELTA Plus XP IRMS (Thermo Scientific, Bremen, Germany) as described previously [35]. Prior to analyses of δ^13^C-DIC, DIC in 1 mL of slurry supernatant was converted to CO_2_ by adding 100 µL phosphoric acid (85%, H_3_PO_4_) overnight at room temperature.

### Nucleic acids extraction, quantification and DNase treatment

The nucleic acids were extracted according to Lueders et al. [36]. Briefly, 2 mL of wet sediment without supernatant was used for cell lysis by bead beating, nucleic acid purification by phenol-chloroform-isoamyl alcohol extraction and precipitation with polyethylene glycol. For the RNA extract, DNA was removed by using the RQ1 DNase kit (Promega, Madison, Wisconsin, USA). DNA and RNA were quantified fluorimetrically using Quant-iT PicoGreen and Quant-iT RiboGreen (both Invitrogen, Eugene, Oregon, USA), respectively.

### Isopycnic centrifugation, gradient fractionation and reverse transcription

Isopycnic centrifugation and gradient fractionation were performed according to the previously described method with modifications [36]. In brief, 600 to 800 ng RNA from each sample was loaded with 240 µL formamide, 6 mL cesium trifluoroacetate solution (CsTFA, GE Healthcare, Buckinghamshire, UK) and gradient buffer solution. The density of the centrifugation medium was adjusted to ∼1.80 g mL^−1^ based on refractive index. RNA was density separated by centrifugation at 124 000 *g* at 20 °C for 65 h using an Optima L-90 XP ultracentrifuge (Beckman Coulter, Brea, California, USA). 13 fractions were obtained from each sample, and RNA in each fraction was precipitated with 1 volume of isopropanol. RNA was quantified and reverse transcription was conducted using the high capacity cDNA reverse transcription kit (Applied Biosystems, Foster City, California, USA).

### Quantitative PCR (qPCR)

Archaeal 16S rRNA and *mcrA* genes were quantified using primer sets 806F/912R and ME2 mod/ME3’Fs 1011 (Table S1), respectively, on a Step One Plus qPCR thermocycler (Applied Biosystems, Foster City, USA); *mcrA* encodes the alpha subunit of methyl coenzyme M reductase, a key enzyme of methanogenic and methanotrophic archaea [37]. Standard curves were based on the 16S rRNA gene of *M. barkeri* and the *mcrA* gene clone A4-67 for archaea and methanogens, respectively. Each 20 µL reaction mixture consisted of 10 µL of MESA Blue qPCR Master Mix (Eurogentec, Seraing, Belgium), 300 nM primers (600 nM for *mcrA* genes), 0.2 mg mL^−1^ bovine serum albumin (Roche, Mannheim, Germany), 1 ng DNA templates or 2 µL of cDNA samples. The qPCR protocol comprised an initial denaturation for 5 min at 95 °C and 40 cycles amplification (95 °C for 30 sec, 58 °C for 30 sec and 72 °C for 40 sec). The detection thresholds were 100–1 000 gene copies with an efficiency of 90–110%.

### Sequencing and bioinformatics analysis

For community composition analysis of single labeling incubations (one substrate labeled, the other unlabeled), “heavy” (1.803–1.823 g mL^−1^, combination of fraction 3, 4, and 5) and “light” (1.777–1.780 g mL^−1^, fraction 11) fractions of RNA-SIP samples were selected according to CsTFA buoyant densities. PCR targeting the V4 region of archaeal 16S rRNA genes was conducted with KAPA HiFi HotStart PCR kit (KAPA Biosystems, Cape Town, South Africa) and barcoded primer sets Arc519F/806R (Table S1). Thermocycling was performed as follows: 95 °C for 3 min; 35 cycles at 98 °C for 20 sec, 61 °C for 15 sec and 72 °C for 15 sec; 72 °C for 1 min. Library construction and sequence read processing were described as previously [34].

### Lipid analysis

Total lipids were extracted from ∼4 g of freeze-dried sediment samples from single labeling incubations (one substrate labeled, the other unlabeled) using a modified Bligh-Dyer protocol [38]. Intact polar archaeal ether lipids were purified by preparative high performance liquid chromatography with fraction collection according to the method by Zhu et al. [39]. The intact archaeal lipid fraction was converted to hydrocarbon derivatives including phytane, phytenes, biphytane, and biphytanes containing cycloalkyl rings (Figure 3C, Figure S1) according to the method by Liu et al. [40]. Archaeal lipid derivatives were quantified by GC-FID (Thermo Scientific, Bremen Germany) and the carbon isotope composition was measured via GC-IRMS on a Thermo Finnigan Trace GC connected with a DELTA V Plus IRMS system via GC Isolink (Thermo Scientific, Bremen Germany). The detailed chromatographic and mass spectrometric parameters were described by Kellermann et al. [41].

### δ^13^C calculation

The proportion of methane from DIC 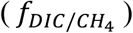 was calculated based on the fractional abundance of ^13^C (^*^13^*^*F*) of methane, methanol (MeOH) and DIC in the incubation with ^13^C-DIC and MeOH. According to a two-end member model, DIC and MeOH are two main carbon sources for methane production expressed as follows:

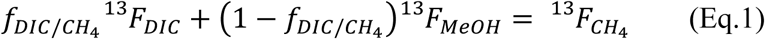

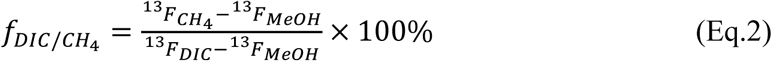

where ^13^*F* is obtained from the δ notation according to F = R/(1+R) and R = (δ/1000+1) * 0.011180 [42]. 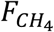 and ^13^*F*_*DIC*_ were the fractional ^13^C abundance of methane and DIC at harvest time, and ^13^*F*_*MeOH*_ that of MeOH in the medium at the start.

^13^C label incorporation ratios from MeOH or DIC in single labeling experiments were calculated from the ^13^C abundance increase relative to the ^13^C label strength via Eq. 3 and Eq. 4. *X*_*MeOH*_ and *X*_*DIC*_ signify the ^13^C incorporation ratio from MeOH and DIC, respectively. 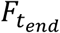 and 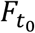 are the ^13^C fractional abundance of lipids harvested at t_end_ and t_0_.

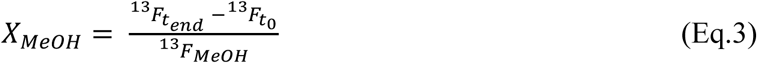

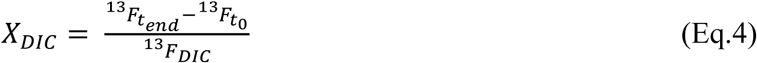

Given that the single labeling incubations were conducted with the same treatment, i.e., 1 mM methanol and 10 mM DIC, the relative proportion of DIC for lipids biosynthesis (*f*_*DIC/lipid*_) was estimated from the ^13^C incorporation ratios (*X*_*MeOH*_ and *X*_*DIC*_) in these single labeling incubations as follow:

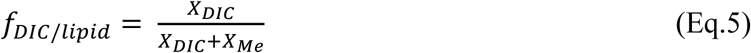

## Results

### Methanogenesis and gene copy numbers in ^13^C-amended sediment slurry incubations

In order to examine carbon labeling into RNA and lipids of methylotrophic methanogens in anoxic marine environments, sediment slurries from two depths amended with or without ^13^C-methanol (1 mM) and ^13^C-DIC (10 mM) were incubated at 10 °C. Samples from SRZ and MZ incubations showed methanogenesis started after 20 and 10 days, respectively (Figure 1A). Amended ^13^C-DIC was diluted into the sediment endogenous DIC pool to about 70–84%, which was more obvious in samples from MZ than SRZ (Table 1). In SRZ sediment incubations with ^13^C-DIC and unlabeled methanol, up to 10.3% of methane originated from ^13^C-DIC (Table 1).

**Figure 1.**
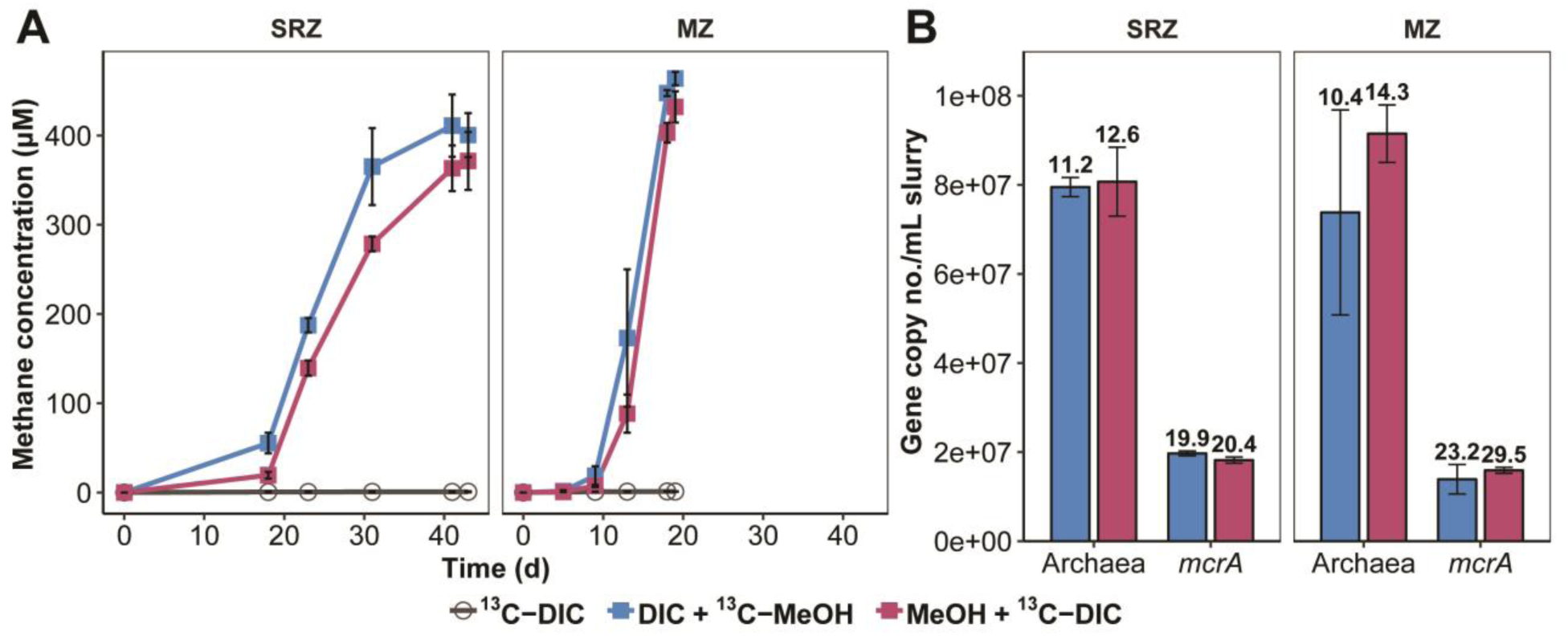
Dynamics of methane formation and archaeal populations in stable isotope probing incubations with SRZ and MZ sediment samples. (A) Methane concentrations in SIP incubations. Methane data is presented as average values (n = 3, error bar = SD). (B) Gene copy numbers of archaea (16S rRNA genes) and methanogens (*mcrA* gene). Gene copies were quantified based on DNA extracts at harvest. Fold increase of gene copies were indicated above each histogram by comparing gene copies on day 0 after preincubation (n = 3, error bar = SD). DIC: dissolved inorganic carbon, i.e., bicarbonate; MeOH: methanol.

**Table 1.** ^13^C fractional abundance and H_2_ partial pressures in SIP incubations. Data is presented as average values (n = 3).

| Sediment | Substrates | $^{13}F_{\text{DIC}}$ (%) | $f_{\text{DIC}/\text{CH}_4}$ (%)<sup>a</sup> | $\text{H}_2$ (Pa)<sup>b</sup> | Incubation time (d) |
| --- | --- | --- | --- | --- | --- |
| SRZ | DIC + $^{13}\text{C}$-MeOH | $3.7 \pm 0.4$ | $89.3 \pm 0.3$ | NA | 43 |
| MZ | DIC + $^{13}\text{C}$-MeOH | $3.0 \pm 0.0$ | $96.4 \pm 0.1$ | NA | 19 |
| SRZ | MeOH + $^{13}\text{C}$-DIC | $83.6 \pm 0.6$ | $10.3 \pm 0.2$ | $0.1 \pm 0.1$ | 43 |
| MZ | MeOH + $^{13}\text{C}$-DIC | $69.8 \pm 0.7$ | $3.4 \pm 0.1$ | $0.3 \pm 0.0$ | 19 |

<sup>a</sup>Methane proportion from DIC ( $f_{\text{DIC}/\text{CH}_4}$ ) in “methanol + $^{13}\text{C}$ -DIC” incubations was based on Eq.2. <sup>b</sup> $\text{H}_2$ partial pressure was measured on day 23 and 16 for incubation SRZ and MZ sediments, respectively. NA, not analyzed.

The dynamics of the archaeal communities in all incubations was tracked by qPCR of archaeal 16S rRNA genes and *mcrA* genes after methanogenesis ceased (Figure 1B). Archaeal and *mcrA* gene copy numbers increased strongly by 10–14 and 19–30 times for all incubations, respectively (Figure 1B).

### Carbon assimilation into RNA and identification of metabolically active archaea

In preliminary sediment incubations, SIP experiments with ^13^C-methanol had shown that RNA could not be labeled to a sufficiently high extent to become detectable in heavy gradient fractions (e.g., >1.803 g mL^−1^) after isopycnic separation of RNA. Contrastingly, methane formation was strong and archaeal and *mcrA* gene copies increased likewise indicating that methylotrophic methanogens were active (Figures 1A and B). Because of mixotrophic assimilation capabilities in methylotrophic methanogens, i.e., utilizing methylated compounds and DIC, a series of SIP slurry experiments were conducted with combinations of methanol and DIC in order to improve the sensitivity of RNA-SIP: double ^13^C-label (methanol + DIC), single ^13^C-label (one of the substrates labeled), both substrates unlabeled, and a ^13^C-DIC control (Figure 2, Figure S2). After density separation of RNA, different degrees of RNA labeling were detected in isotopically heavy gradient fractions, e.g., >1.803 g mL^−1^ (Figure 2). Strongest ^13^C-labeling, as indicated by largest amounts of RNA found in gradient fractions > 1.803 g mL^−1^, was detected in RNA from incubations with double ^13^C-labeling (Figure 2A), followed by single-label incubations with ^13^C-DIC. For single-label ^13^C-methanol incubations, however, RNA fraction shifts according to density were minor compared to unlabeled incubations, indicating limited assimilation of ^13^C-methanol into RNA of methylotrophic methanogens.

**Figure 2.**
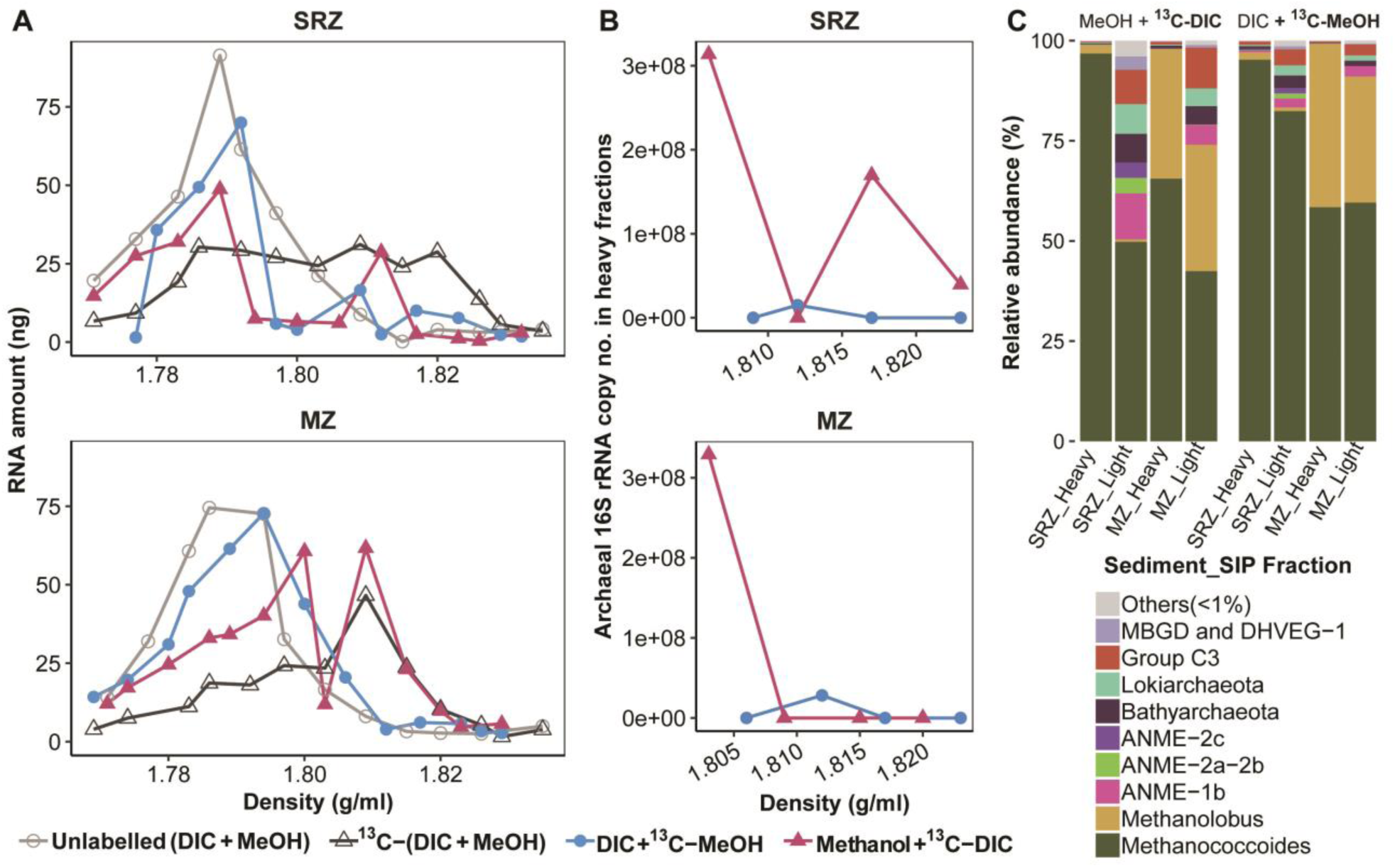
Density distribution of RNA, gene copy numbers, and community composition from SIP incubations with SRZ and MZ sediment after isopycnic separation. (A) RNA profiles from different RNA-SIP experiments. (B) Gene copy numbers of archaeal cDNA in heavy fractions (1.803 g mL^−1^ to 1.823 g mL^−1^) from RNA-SIP experiments. (C) Relative abundances of density separated archaeal 16S rRNA from single-labeling incubations in light (1.771 g mL^−1^ to 1.800 g mL^−1^) and heavy (1.803 g mL^−1^ to 1.835 g mL^−1^) gradient fractions.

In order to estimate ^13^C-labeling levels of methanogens in single SIP experiments (^13^C-methanol or ^13^C-DIC), a series of molecular techniques were applied including qPCR of cDNA in heavy fractions of RNA-SIP samples, archaeal 16S rRNA sequencing from RNA-SIP fractions and δ^13^C value determination of methanogen lipids, e.g., phytanes derived from intact polar archaeol-based molecules. In incubations amended with ^13^C-DIC and unlabeled methanol, archaeal 16S rRNA gene copies were one order of magnitude higher in the heavier gradient fractions (i.e., 1.803–1.823 g mL^−1^) than that in ^13^C-methanol incubations (Figure 2B). Correspondingly, Illumina sequencing of RNA revealed that sequences identified as related to the genera *Methanococcoides* and *Methanolobus* were more dominant in the heavy than in the light fractions from incubations amended with unlabeled methanol and ^13^C-DIC (Figure 2C, S3). Light fractions were overall mainly composed of anaerobic methanotrophic (ANME) archaea, Bathyarchaeota and Lokiarchaeota except for methylotrophic methanogens (Figure 2C). For ^13^C-DIC control incubations, abundances of methanogens were low in SIP samples (Figure S3). For unlabeled methanol incubations, given the low amount of labeled RNA in heavy fractions, no amplicons were obtained, but light fractions showed a high abundance of methylotrophic methanogens (Figure S3). Classifications were confirmed by phylogenetic clustering of cloned 16S rRNA gene fragments (about 800 base pairs) with OTU sequences representing *Methanococcoides* and *Methanolobus* spp. (Figure S4). Sequences of these methanogens accounted for more than 97% of total archaea in heavy gradient fractions. However, known hydrogenotrophic methanogens were undetectable (Figure 2C) although 5 and 10% of methane was formed from DIC in incubations with MZ and SRZ sediments, respectively (Table 1).

In parallel to RNA-SIP, lipid-SIP incubations with SRZ sediment slurries demonstrated δ^13^C values of phytane and phytenes being more positive in ^13^C-DIC and unlabeled methanol treatment than that in ^13^C-methanol amendments, while the opposite was found in MZ sediment incubations (Figure 3A). After elimination of ^13^C-DIC dilution effects by ambient inorganic carbon, DIC contributions to lipids ranged from 59.3% to 86.1% in SRZ sediment incubations, which was constantly higher than that of MZ sediment incubations (52.7% to 56.4%).

**Figure 3.**
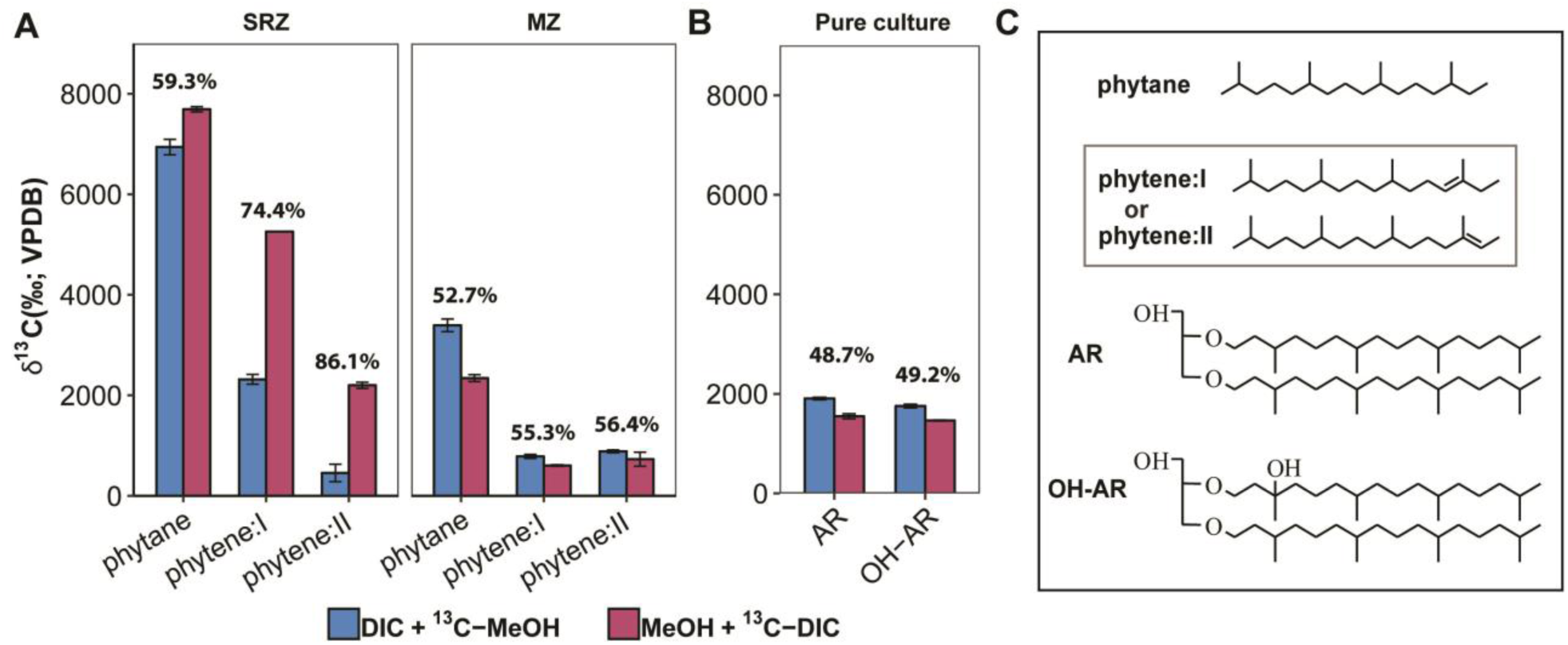
Lipid-SIP experiments from sediment incubations (natural community) and pure cultures in Widdel medium. Lipid δ^13^C values were measured in homogenized samples after methanogenesis had ceased. (A) δ^13^C values of phytanes in sediment incubations with 70% ^13^C-DIC. Phytane originates from intact polar archaeol lipids and phytenes (phytene I and phytene II) derive from intact polar hydroxyarchaeol lipids. *f*_DIC/lipid_ is indicated on the top of bars from single labeling incubations based on Eq.5. (B) δ^13^C values of archaeol (AR) and hydroxyarchaeol (AR-OH) in pure culture of *M. methylutens* treated with 5% ^13^C-labeled substrates (methanol or DIC). (C) Structures of archaeal lipids. Enclosed structures of phytenes in Figure C were tentatively assigned according to GC-MS mass spectra (Figure S6) [71]. Data is expressed as average values (n = 3, error bar=SD).

To understand how carbon is assimilated into lipids by methylotrophic methanogens, pure culture incubations of *M. methylutens* were performed with 5% of the ^13^C-labeled substrates (i.e., DIC or MeOH) and the dominating archaeal lipids archaeol (AR) and hydroxyarchaeol (OH-AR) were directly analyzed without cleavage. In contrast to sediment incubations, lipids showed lower *f*_DIC/lipid_ (∼49%) based on the carbon incorporation in single labeling incubations (Figure 3B).

### Methylotrophic methanogenesis from DIC in autoclaved slurry incubations

The unexpectedly high proportion of methane formed from DIC in methanol amended sediment slurry incubations (Table 1) prompted us to investigate the underlying mechanism in more detail. Thus, autoclaved sediment slurries were used as a surrogate of natural sediment, but with all microorganisms killed, and inoculated with the obligate methylotroph *M. methylutens*. Hematite and magnetite known to serve as electron acceptors or conductors [43,44] were added along with humic acid, and AQDS as electron shuttles, as well as an additional electron donor (H_2_), which are all known to stimulate methanogenesis [43,44,45,46].

Methane concentrations in incubations with hematite and humic acid were higher than that of the other incubations after 7 days (Figure 4A). Although methane production rates were low, methane proportions from DIC in treatments with *M. methylutens* alone, H_2_, AQDS and magnetite were much higher (*f*_*DIC/CH4*_, ∼10%) than that in incubations with hematite and humic acid (∼2%) (Figure 4B). Linear regression showed a strong correlation between methane production rate and CO_2_-dependent methanogenesis by methylotrophic methanogens on day 3 and 5 of the incubations, indicating that lower methanogenesis rates triggered higher levels of methane formation derived from ^13^C-DIC (Figure 4C).

**Figure 4.**
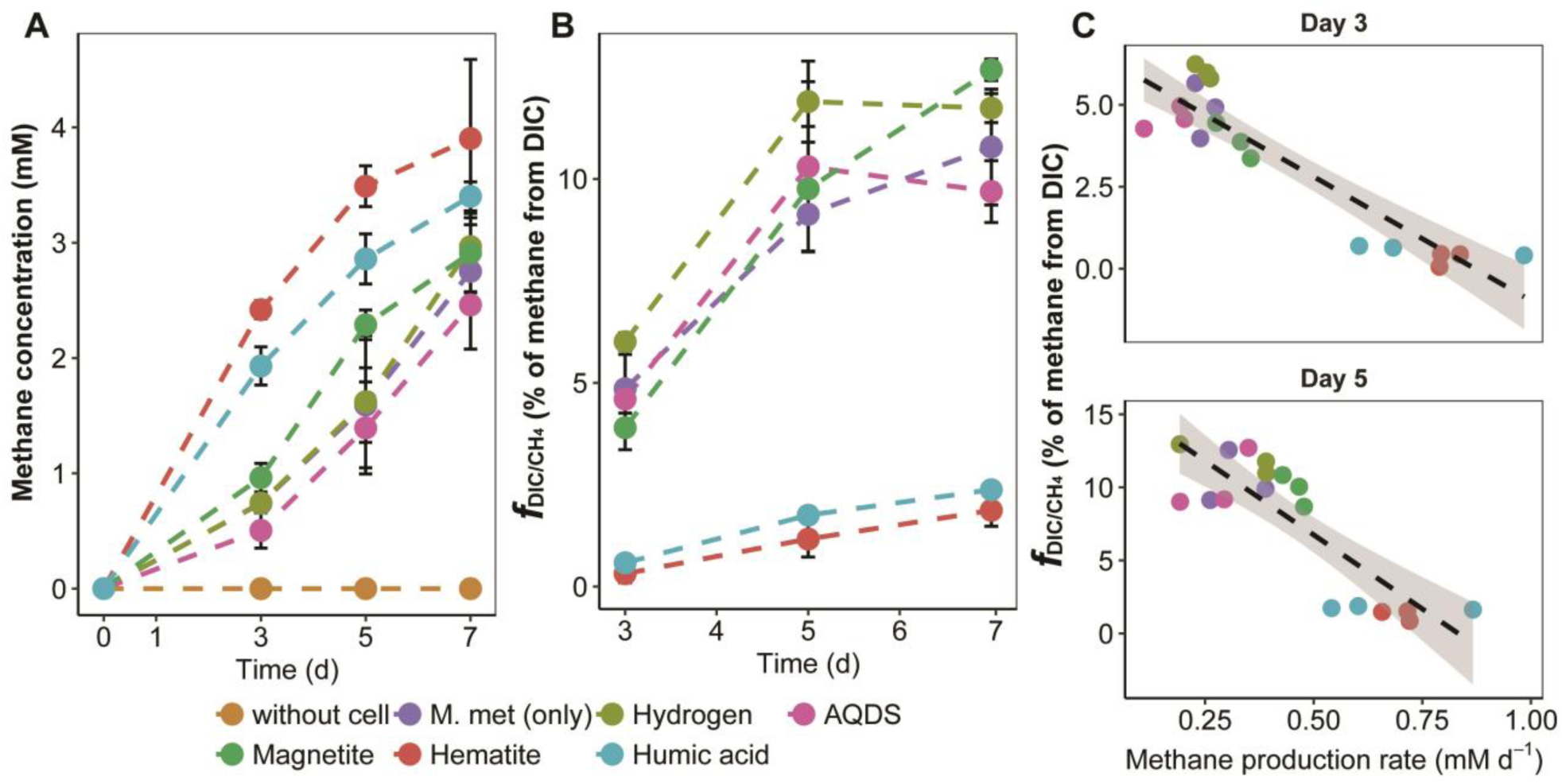
Methane production from DIC during methylotrophic methanogenesis in autoclaved slurry supplemented with pure culture of *M. methylutens*. (A) Total methane concentrations in headspace. (B) Proportion of methane derived from DIC. Methane proportion from DIC 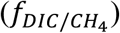 was calculated according to Eq.2. Data is expressed as average values (n = 3, error bar = SD). (C) Linear correlation between methanogenesis rate and 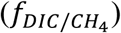 after 3 and 5 days. Day 3: Pearson?s *r* = −0.92, *P* < 0.001, C1(0.95) = −0.79 > *r* > −0.97; Day 5: Pearson?s *r* = −0.85, *P* < 0.001, C1(0.95) = −0.62 > *r* > −0.94. M. met: *M. methylutens*.

## Discussion

In this study, we utilized RNA-SIP employing ^13^C-DIC and methanol and successfully identified methylotrophic methanogens in both, SRZ and MZ sediments of the Helgoland mud area in the North Sea. We demonstrated that the addition of ^13^C-DIC is necessary to detect label in RNA of methylotrophic methanogens rather than using ^13^C-methanol as energy substrate alone. We further evaluated carbon utilization patterns of the methylotrophic methanogens by lipid-SIP and identified an unexpectedly high DIC assimilation into characteristic lipids within SRZ sediment. Isotope probing experiments unexpectedly revealed that up to 12% of methane was formed from DIC by the presumably “obligate” methylotrophic methanogen, *M. methylutens*, thereby suggesting an explanation for the elevated DIC incorporation into biomass.

### Carbon assimilation by methylotrophic methanogens in sediment incubations

Nucleic acids-SIP techniques depend on ^13^C-labeling levels of DNA or RNA molecules, from which carbon assimilation can be reconstructed and compared to the known pathway of nucleic acid biosynthesis from methyl-groups in methanogens [47,48,49,50,51,52]. The current pathways show that only one carbon atom stems from methanol in ribose-5-phosphate while 25% to 40% of carbon in nucleobases originates from the methyl carbon of the substrate (Figure 5). This is corroborated by our RNA-SIP experiments using ^13^C-labeled methanol alone, but RNA was not found to be labeled effectively enough for density separation and further sequence analysis. However, by additionally using ^13^C-DIC, we found high 16S rRNA copy numbers (Figure 2B) and a high representation of known methylotrophic methanogens (Figure 2C) in the heavy RNA gradient fractions, successfully recovering ^13^C-labeled RNA of methylotrophic methanogens in the SRZ and MZ sediments of the Helgoland mud area. Consequently, RNA labeling will be more effective in methylotrophic methanogenic archaea by DIC than methanol. Combined with downstream analysis including qPCR, 16S rRNA sequencing and cloning, we directly show that members of the genus *Methanococcoides* were the predominantly active methylotrophic methanogens in SRZ incubations, while *Methanococcoides* together with *Methanolobus* were dominating in MZ incubations. Hence, addition of ^13^C-labeled DIC or a combination of both substrates labeled enables tracking of methylotrophic methanogens via RNA-SIP techniques. For carbon assimilation into nucleic acids of these methanogens, both proposed biosynthesis pathway of nucleic acid and labeling strategy of RNA-SIP confirmed inorganic carbon as the main carbon source for nucleic acids.

**Figure 5.**
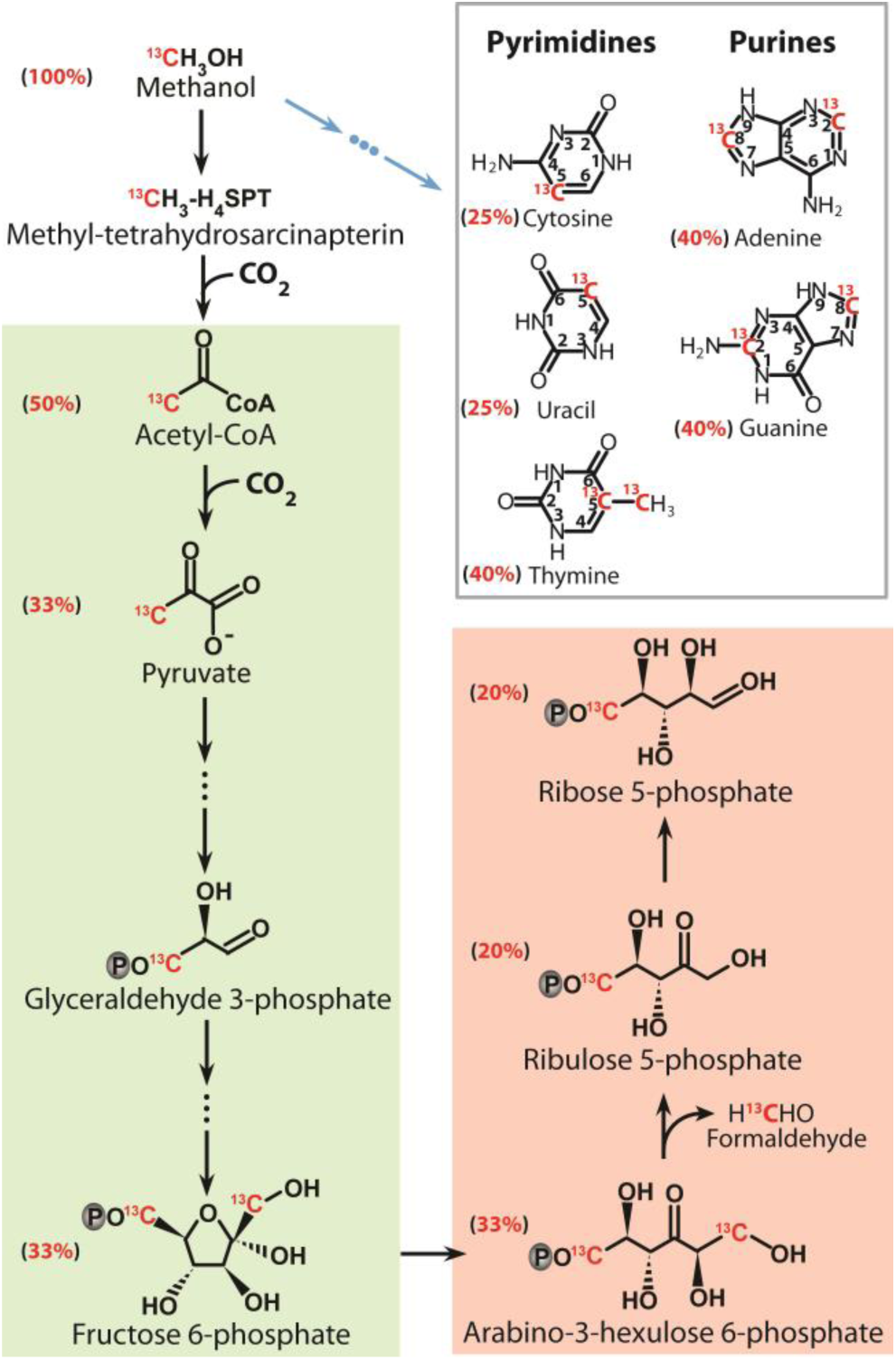
Carbon assimilation into nucleic acids in methylotrophic methanogens. Biosynthesis of nucleotide moieties, the pyrimidine and purine bases, as well as the C5-carbon from ^13^C-labeled methanol in methylotrophic methanogens based on previous studies [47,48,49,50,51,52] with final carbon contribution from methanol added besides the compounds. Black arrows indicate ribose synthesis and the blue arrows represent synthesis of base moieties in nucleosides. The reverse gluconeogenesis pathway is displayed in green and the reverse ribulose monophosphate pathway in pink.

Since carbon assimilation into biomass cannot be quantified by RNA-SIP, lipid-SIP was used. In lipid-SIP analysis, we evaluated ^13^C-incorporation into intact polar archaeol- and hydroxyarchaeol diether molecules, which are the dominant lipids produced by moderately thermophilic methanogenic archaea [53,54,55], via phytane and phytene side-chain analysis (Figure 3). These moieties were the only ones being ^13^C-labeled while tetraether-derived biphytane and cycloalkylated biphytanes as indicators of archaea such as Thaumarchaeota [56], anaerobic methanotrophs [57,58] or Bathyarchaeota [59] did not show a ^13^C incorporation (Figure S5). This was corroborated by our sequencing results demonstrating that methylotrophic methanogens were the dominant archaea in the heavy fractions and that the relative abundances of other archaea were very low or even below detection (Figure 2C).

By evaluating ^13^C incorporation into methanogen-derived phytane and phytenes, lipid-SIP provides insight into methanogen activities and carbon utilization. As the main precursors of ether lipids in archaea, biosynthesis of isopentenyl diphosphate (IPP) and dimethylallyl diphosphate (DMAPP) proceeds via the modified mevalonate pathway [60,61,62]. In this pathway, mevalonate-5-phosphate is decarboxylated to IPP, in which three out of five carbon atoms are derived from methanol (Figure 6). DMAPP is further converted to geranylgeranyl diphosphate (GGPP), which receives 60% of its carbon from methanol, suggestive of a lower DIC contribution to isoprenoid chains than methanol. This was supported by the fact that archaeol and hydroxyarchaeol contained more methanol-derived than DIC-derived carbon using a pure culture of *M. methylutens* (Figure 3B). However, unlike the proposed lipid biosynthesis pathway and the pure culture, clearly more DIC was assimilated into lipids than methanol in both sediment incubations, which was most prominent in the sediment from the SRZ (Figure 3A). We, moreover, detected that ∼10% of methane produced was derived from DIC during the SIP experiments and using *M. methylutens* in autoclaved sediment slurry incubations (Table 1, Figure 4). From the lipid biosynthesis pathway it is very likely that part of the DIC is converted to methyl-tetrahydrosarcinapterin (CH_3_-H_4_SPT) and further reduced to methane, and thus, such CH_3_-H_4_SPT generated from CO_2_ will be simultaneously available for lipid biosynthesis (Figure 6) leading to the ^13^C-enrichment of the lipid pool.

**Figure 6.**
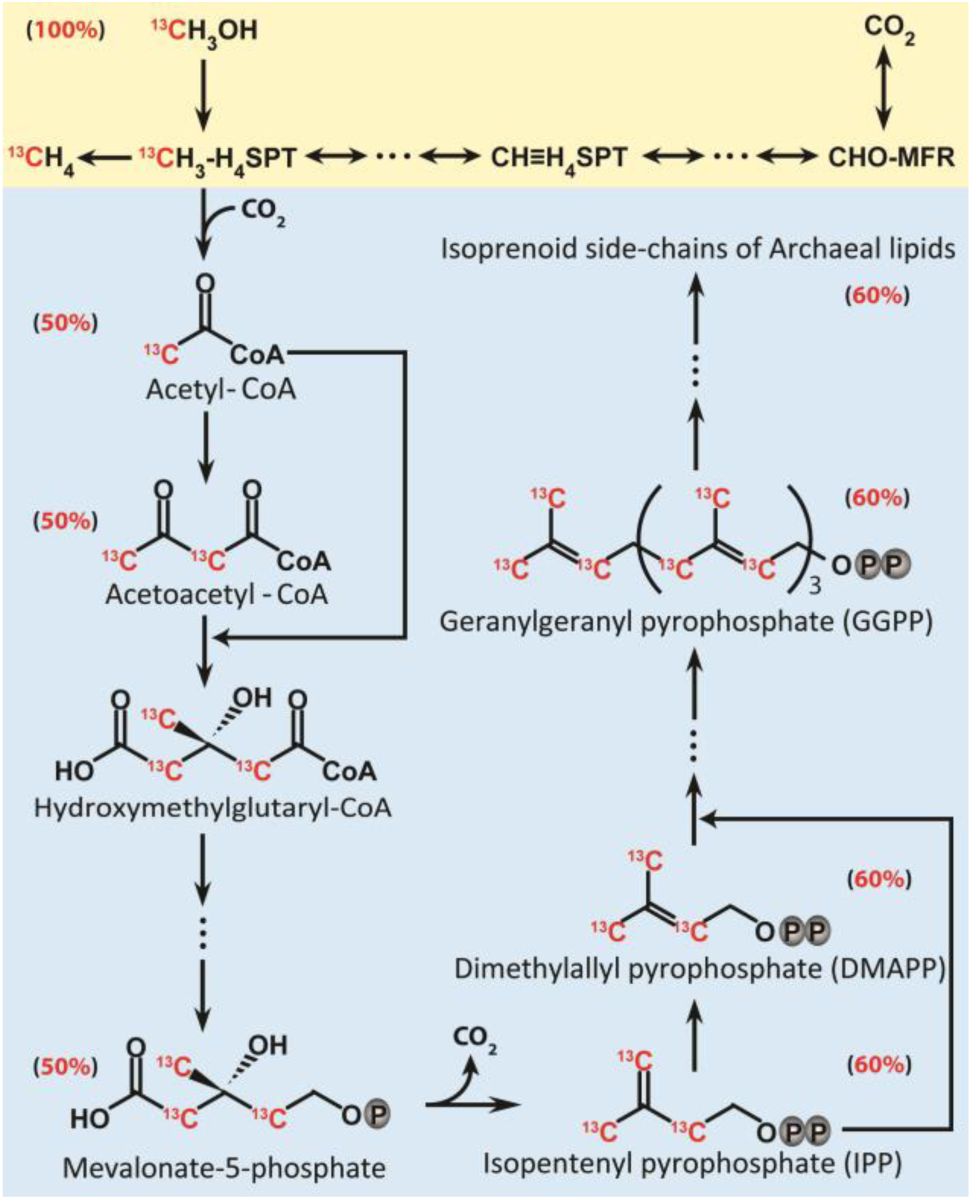
Carbon assimilation into lipids in methylotrophic methanogens. Methylotrophic methanogenesis pathway from methanol (yellow) and carbon assimilation pattern into isoprenoid chains of archaeal lipids (blue) with carbon contribution from ^13^C-methanol added besides the compounds. The pathway of archaeal lipid biosynthesis is based on previous studies [60, 72,73].

### CO_2_ reduction to methane by obligate methylotrophic methanogens

There are two types of CO_2_-dependent methanogenesis: i) Hydrogenotrophic methanogenesis by the orders of *Methanopyrales, Methanococcales, Methanobacteriales, Methanomicrobiales, Methanocellales* and *Methanosarcinales* [2,3]. These methanogens contain F_420_-reducing [NiFe]-hydrogenase to catalyze F_420_ reduction by H_2_ [63]. ii) Mediation by interspecies electron transfer between bacteria and some members of the *Methanosarcinales*. CO_2_ reduction to methane was observed in *Methanosaeta* and *Methanosarcina* during syntrophic growth with *Geobacter* species on alcohols (ethanol, propanol, and butanol), as electrons generated from *Geobacter* are directly transferred to methanogens to reduce CO_2_ [64,65,66,67].

Based on our SIP incubations with SRZ sediment showing 10% of methane generation from DIC at low H_2_ partial pressure (< 0.3 Pa) (Table 1) and the overall lack of hydrogenotrophic methanogens in RNA-SIP fractions (Figure 2C), we argue that H_2_-dependent methanogenesis does not play a role [2]. Similarly, in autoclaved sediment slurry (Figure 4B), the “obligate” methylotroph *M. methylutens* generated methane from CO_2_ without a hydrogen (or electron) supplying partner microorganism.

Members of the genus *Methanococcoides* are considered as obligate methylotrophic methanogens since no F_420_-reducing [NiFe]-hydrogenase was detected in their genomes [60, 68], ruling out hydrogenotrophic methanogenesis in sediment incubations. Nevertheless, part of the methane formed during methylotrophic methanogenesis by *M. methylutens* was from CO_2_, especially when methane production rates were low (Figure 4C). Apparently, at high rates of methanol dissimilation to CO_2_, the reverse pathway of CO_2_ reduction to methane was outcompeted. Higher rates of methylotrophic methanogenesis can be achieved potentially by amendments in autoclaved slurries using hydrogen as electron donor, electron conductors (hematite, magnetite) and electron shuttles (humic acid, AQDS); these compounds are known to affect electron utilizations in microorganisms and stimulate methanogenesis [43,44,45,46]. In our incubations, we found humic acid and hematite most strongly stimulating methylotrophic methanogenesis. Although the underlying mechanism is beyond the scope of the current paper, methylotrophic methanogens in our incubations could take advantage of hematite as potential electron conductor [43,44] or humic acid as electron shuttle [46] as indicated by a higher rate of methanogenesis compared to the other treatments (Figure 4A).

It has been shown that 3% of methane was produced from CO_2_ during methylotrophic methanogenesis of *Methanosarcina barkeri* (i.e., a facultative methylotroph) without the addition of H_2_ [20], which is similar to about 2.5% of methane generated from CO_2_ by *M. methylutens* (i.e., a presumed “obligate” methylotroph) in our study (Table S2). However, methane generation from CO_2_ increased to 10% in SRZ sediment incubations (Table 1), highlighting the importance of the activity of concomitant CO_2_ reduction during methylotrophic methanogenesis in marine sediments. It seems that the substantially higher methanogenesis rates in pure cultures decrease the amount of methane produced from CO_2_. In marine sediment methylotrophic methanogenesis rates are likely lower than those in pure cultures because of the limitation in methylated substrates [6, 14], strongly suggesting that methane generation from CO_2_ by methylotrophic methanogens is underestimated under *in situ* conditions. Thus, CO_2_ conversion to methane has to be considered when estimates of *in situ* methylotrophic methanogenesis in marine sediments are performed.

### Conclusions

In this study, we have shown that ^13^C-DIC is required as co-substrate for successful identification of methylotrophic methanogens by RNA-SIP in marine sediments. DIC is the main carbon source for biosynthesis of nucleic acids in these methanogens and thus using ^13^C-methanol as energy and carbon substrate alone is insufficient in SIP experiments. Given the intricacies of known assimilatory pathways in methanogenic archaea as a functional group, it might be necessary to at least check for the possibility of DIC as a main assimilatory carbon component in all methanogens for successful SIP experiments. In general, it seems that archaea have a propensity for using DIC as a carbon source for assimilation [59, 69], possibly as an evolutionary adaptation to environments with limited availability of organic carbon [70]; in our laboratory, we are currently unveiling more DIC assimilating microorganisms in marine sediments.

But beyond known pathways, we detected an unexpectedly high amount of methane (> 10%) formed from DIC. This finding strongly suggests that the alleged obligate methylotroph studied here was rather mixotrophically converting both available substrates (DIC, methanol) to methane. This seems to be a conundrum, however, our detailed labeling studies showed that the kinetics of substrate utilization apparently is a decisive factor in channeling more or less CO_2_ into the pathway of methanogenesis: more methane formed from CO_2_ when the overall kinetics were slow, and vice versa. It is not obvious currently, what the benefit for the cell is, since more methane could be formed when methanogenesis proceeds by disproportionation of methanol. From an ecological perspective, DIC is a much more pertinent substrate than methanol (or other methyl compounds) in marine sediments [4, 6], and thus, we speculate that more DIC reduction by obligate methylotrophic methanogens occurs *in situ* than is currently known. A larger proportion of DIC-dependent methanogenesis from methyl compounds should also impact carbon fractionation, and thus, delta ^13^CH_4_ might be overprinted by such mixotrophic methanogenesis. Thus, the CO_2_ reduction to methane and assimilation into biomass by obligate methylotrophic methanogens plays a much more important role in the environment than was previously known.

## Acknowledgments

This study was supported by the Research Center/Cluster of Excellence "The Ocean in the Earth System? (MARUM) funded by the Deutsche Forschungsgemeinschaft (DFG) and by the University of Bremen. Xiuran Yin and Weichao Wu were funded by the scholarship from China Scholarship Council (CSC). We thank the captain, crew and scientists of R/V HEINCKE expeditions HE443.

## Conflict of interest

The authors declare that they have no conflict of interest.

## Data availability

Sequencing data have been submitted to GenBank Short Reads Archive with accession numbers from SRR8207425 to SRR8207442.

