## Supplementary materials for "CO_2_ conversion to methane and biomass in obligate methylotrophic methanogens in marine sediments"

Running title: Methane formation from CO_2_ in *Methanococcoides*

Correspondence:

Michael W. Friedrich,

Microbial Ecophysiology Group, Faculty of Biology/Chemistry, University of Bremen, PO

Box 33 04 40, D-28334 Bremen, Germany

* These authors contributed equally to this work.

^‡^ Current address: Department of Biogeochemistry of Agroecosystems, University of Goettingen, Goettingen, Germany

**Clone library construction**

A clone library of archaeal 16S rRNA gene fragments (~800 bp) was constructed to confirm the accuracy of classification by using short Illumina sequences (143 base pairs). PCR was conducted with primer set of 109F/912R (Table S1) and ALLin RPH polymerase Kit (highQu, Kraichtal, Germany) according to the protocol of the manufacturer. The template cDNA was used from the heavy fractions of RNA-SIP sample of the MZ incubations amended with ^13^C-DIC and unlabeled methanol. Thermocycling was performed as follows: 95 °C for 3 min; 40 cycles at 95 °C for 30 sec, 58 °C for 45 sec and 72 °C for 45 sec; 72 °C for 10 min. Purified PCR products were ligated into the pGEM-T vector (Promega, Mannheim, Germany) and transformed into *Escherichia coli* JM109 competent cells (Promega, Mannheim, Germany) according to the manufacturer. White colonies were randomly picked and cell material directly subjected to colony PCR with the following cycling parameters: 95 °C for 5 min; 28 cycles at 95 °C for 30 sec, 55 °C for 45 sec and 72 °C for 1 min; 72 °C for 5 min. Amplicons of 8 clones were submitted to LGC Genomics (Berlin, Germany) for Sanger sequencing. Sequences have been deposited at GenBank with accession numbers from MK434328 to MK434335.

**Table S1.** Primers used in this study

| **Target gene** | **Primer** | **Reference** |
| --- | --- | --- |
| Archaeal 16S rRNA gene | 806F  (5’-ATTAGATACCCSBGTAGTCC-3’) | [1] |
| Archaeal 16S rRNA gene | 912R  (5’-GTGCTCCCCCGCCAATTCCTTTA-3’) | [2] |
| *mcr*A | ME2 mod  (5’-TCATBGCRTAGTTNGGRTAGT-3’) | [3] |
| *mcr*A | ME3’Fs  (5’-GTCNGGTGGHGTMGGSTTYAC -3’) | [4] |
| Archaeal 16S rRNA gene | Arch519F  (5’-CAGCMGCCGCGGTAA-3’) | [5] |
| Archaeal 16S rRNA gene | Arch806R  (5’-GGACTACVSGGGTATCTAAT-3’) | [6] |
| Archaeal 16S rRNA gene | 109F  (5’-ACKGCTCAGTAACACGT-3’) | [7] |

**Table S2.** Methanogenesis from methanol and DIC in pure culture of *M. methylutens* grown in Widdel medium. Data is expressed as average values (n = 3).

| Substrates | δ^13^C-methane,  (‰; VPDB) | δ^13^C-DIC_day 0, (‰; VPDB) | δ^13^C-DIC_day 11, (‰; VPDB) | Methane from labeled substrate, % |
| --- | --- | --- | --- | --- |
| DIC + 5% ^13^C-MeOH | 4620 ± 160 | NA | NA | 97.1 ± 3.2 |
| MeOH + 5% ^13^C-DIC | 63.8 ± 5.5 | 4170 ± 80 | 3520 ± 82.0 | 2.3 ± 0.1 ~ 2.6 ± 0.2 |

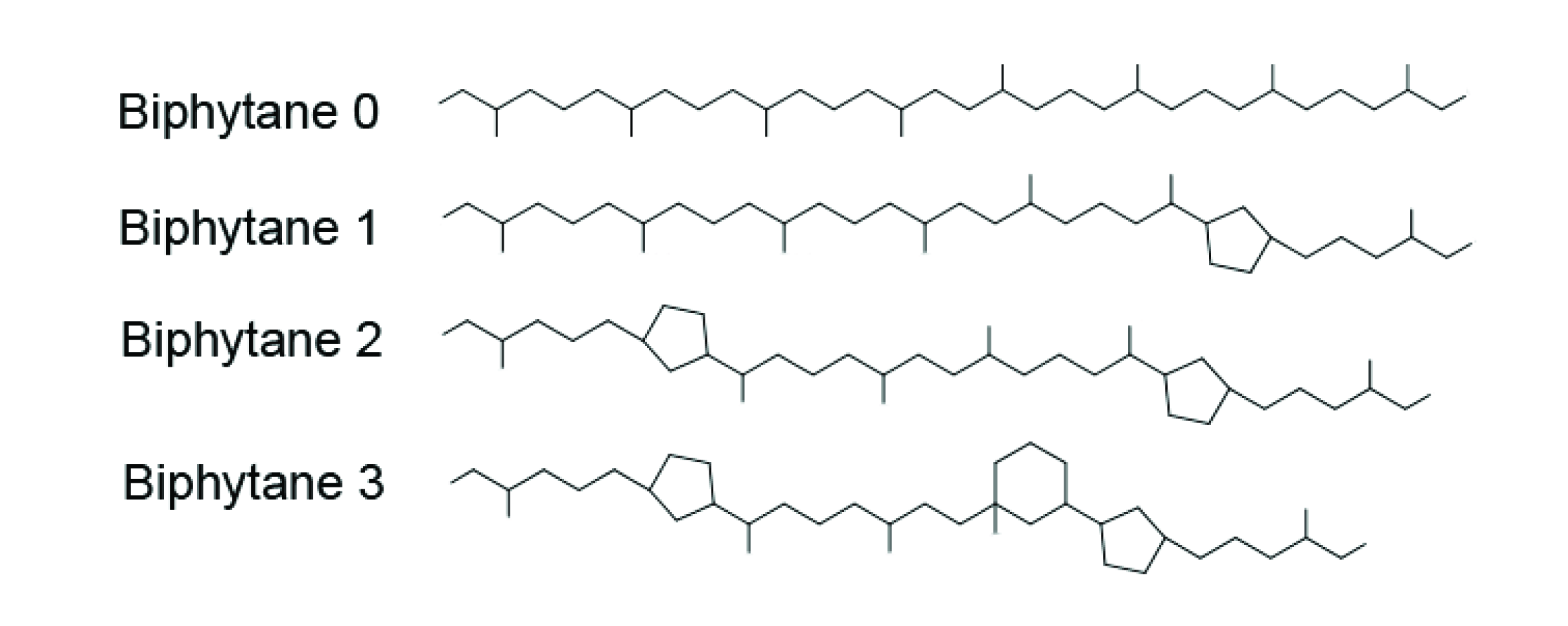

**Figure S1.** Structures of biphytane moieties released from intact polar glycerol diphytanoyl glycerol tetraether fraction.

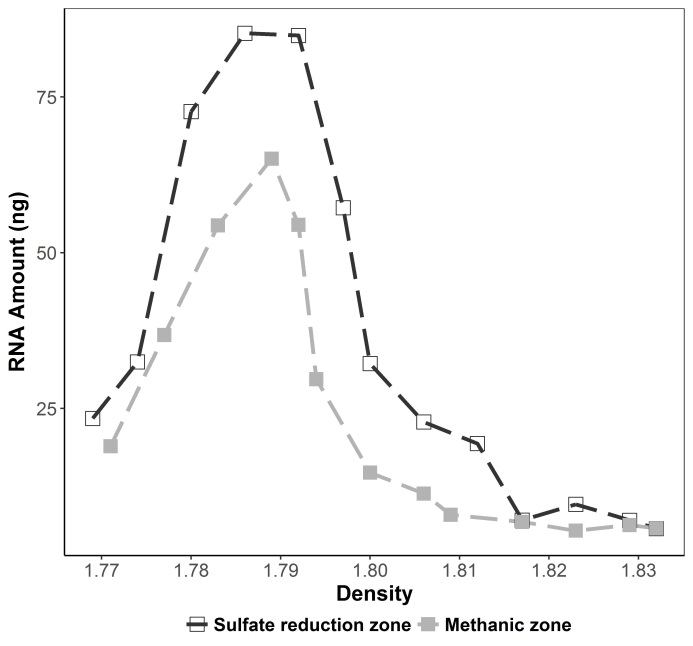

**Figure S2.** RNA-SIP profiles from slurry incubations amended with ^13^C-DIC only (no methanol added). Samples were harvested in parallel to methanol amended incubations.

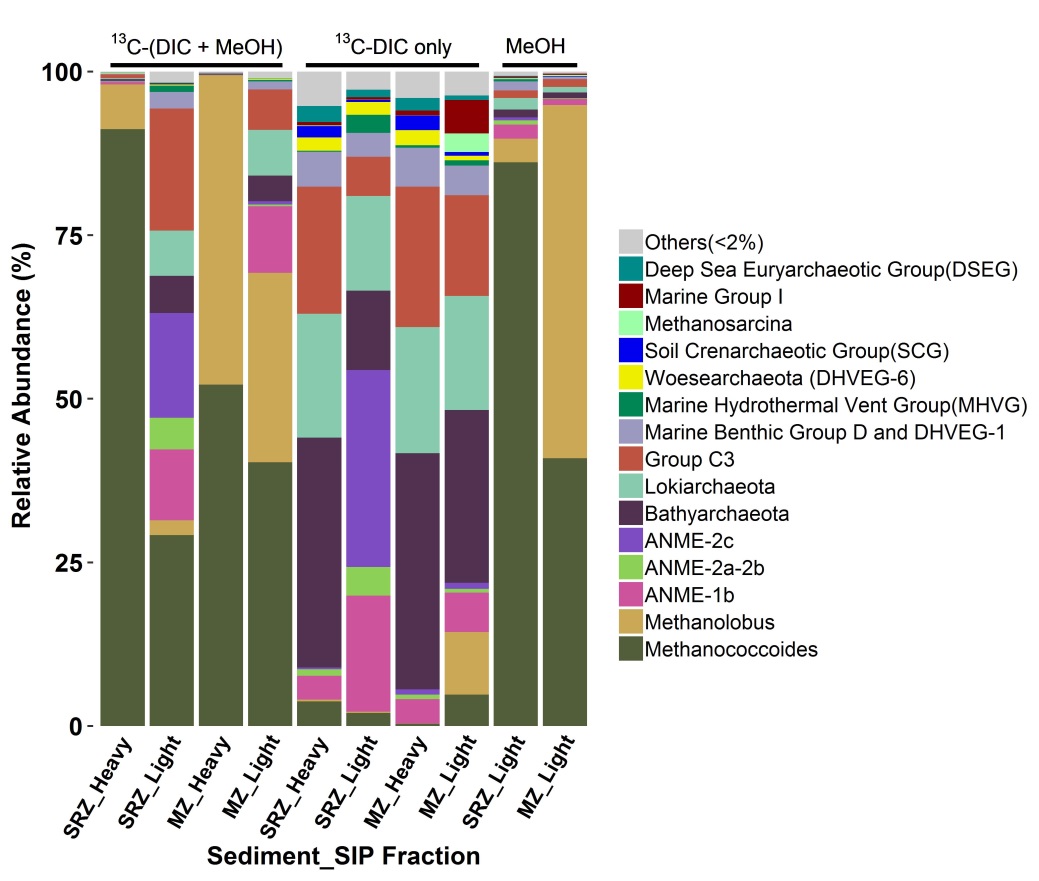

**Figure S3.** Relative abundance of archaeal 16S rRNA in the RNA-SIP samples from double-labeling incubations (^13^C-DIC + ^13^C-methanol) and control incubations (^13^C-DIC or unlabeled-methanol). No data are shown for the unlabeled-methanol controls from the SRZ and MZ due to low amount of RNA in the heavy fraction.

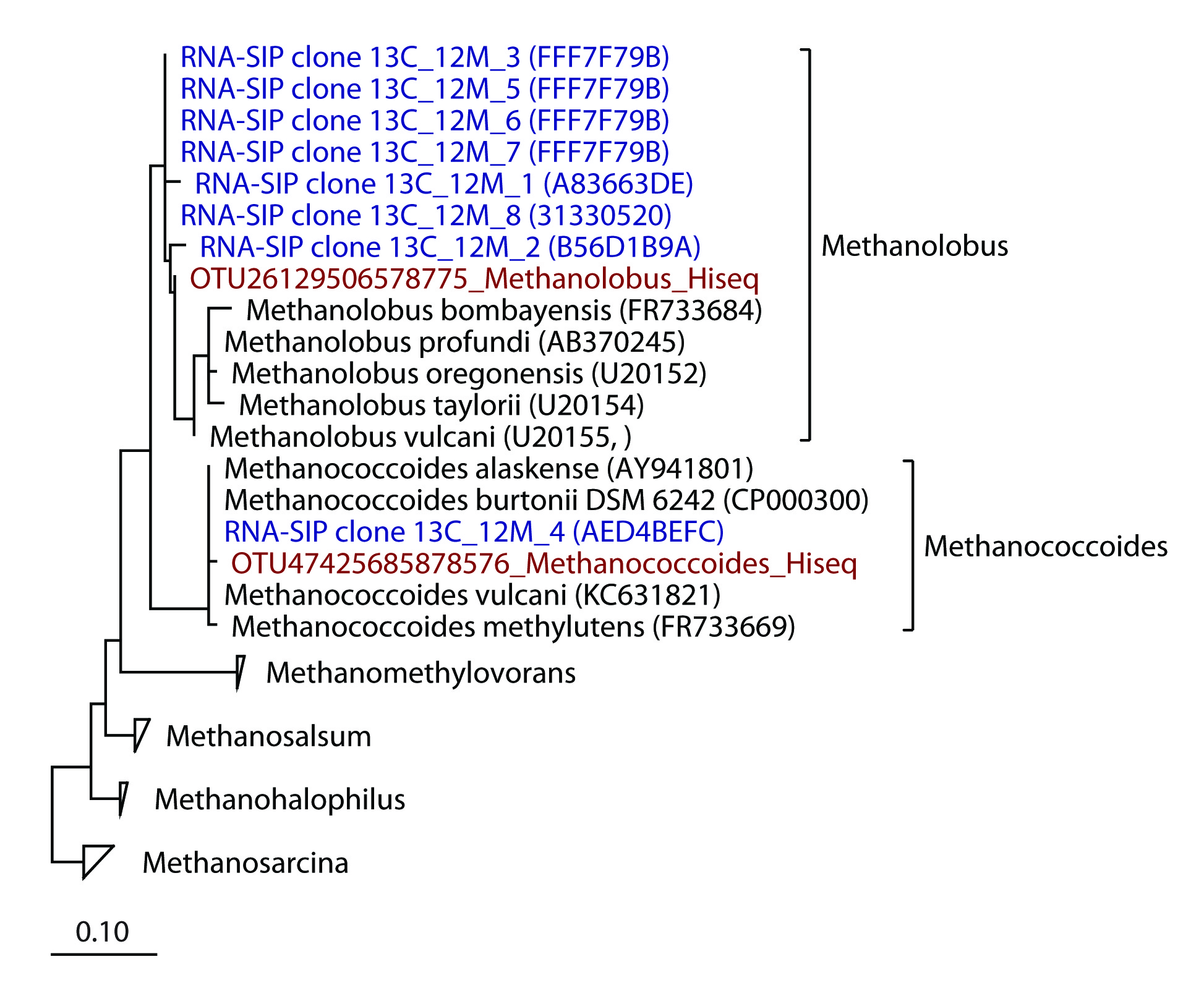

**Figure S4.** Phylogenetic tree of 16S rRNA genes from clone library (blue) and Illumina sequencing (red). Clone sequences were assembled by using SeqMan software (Version 8.0.2) and aligned online by Silva aligner (<https://www.arb-silva.de/aligner/>). The aligned sequences were input into ARB (Version 6.0.2). Aligned clone sequences and know sequences of *Methanosarcinaceae* in SILVA SSURef database (Release 132) were selected to build a phylogenetic tree using maximum likelihood algorithm and bootstrapping (n=1000). The two dominant OTUs of *Methanosarcinaceae* from Hiseq Illumina sequencing (red) were aligned and added to the tree using the ARB parismony tool.

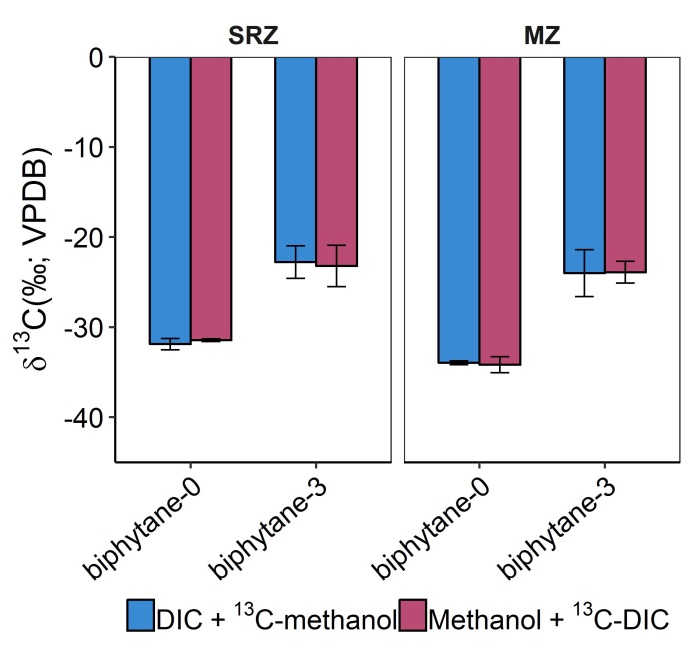

**Figure S5.** δ^13^C values of biphytanes released from the intact ploar glycerol diphytanoyl glycerol tetraether fraction. Biphytane-1 and biphytane-2 were undetectable. Determiantion of carbon isotope values was performed after methanogenesis had ceased. Data is expressed as average values (n = 3, error bar = SD).

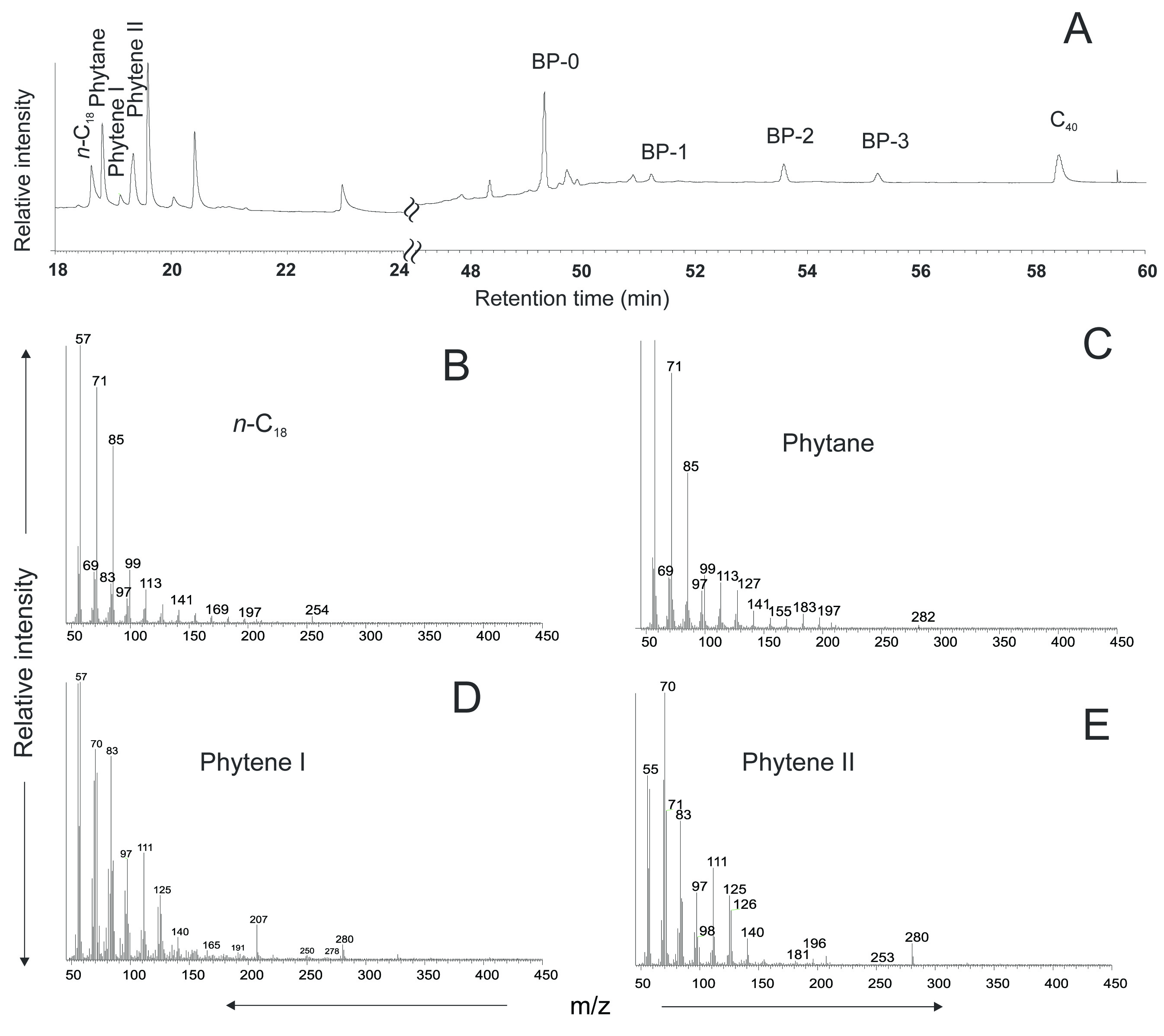

**Figure S6.** Chromatogram of phytane and phytenes released from intact polar lipid fraction in the incubations of the MZ sample amended with ^13^C-DIC and unlabeld methanol (A) and the mass spectra of compunds in the retention of interests (B to E).
